# Structural basis for site-specific histone H3 acetylation-dependent regulation of RNAPII transcription through nucleosomes

**DOI:** 10.64898/2026.05.02.722397

**Authors:** Takumi Oishi, Suguru Hatazawa, Tomoya Kujirai, Sae Suzuki, Koki Nakatsu, Gosuke Hayashi, Junko Kato, Yuki Kobayashi, Mitsuo Ogasawara, Haruhiko Ehara, Shun-ichi Sekine, Yoshimasa Takizawa, Hitoshi Kurumizaka

## Abstract

Acetylation of histone H3 regulates chromatin dynamics and transcription, but how specific acetylation sites affect RNA polymerase II (RNAPII) transcription through nucleosomes remains unclear. Here, we found that H3 acetylation at Lys56 and Lys122 markedly enhances RNAPII transcription through nucleosomes, whereas acetylation at Lys64 has little effect. To elucidate the structural basis for these functional differences, we determined cryo-electron microscopy (cryo-EM) structures of nucleosomes bearing site-specific acetylation at H3K56, H3K64, or H3K122. The cryo-EM structures revealed that H3K56ac and H3K122ac locally weaken histone-DNA interactions at the DNA entry/exit region and near the dyad, respectively, while H3K64ac induces no detectable structural changes. These structural differences correlate with the observed transcriptional outcomes, indicating that acetylation at H3K56 and H3K122, but not H3K64, alleviates the nucleosomal barrier to RNAPII progression. Our findings provide a model in which specific acetylations within the histone fold domain of H3 fine-tune nucleosome dynamics to facilitate RNAPII transcription.

## Introduction

Chromatin plays a central role in regulating access to genomic DNA during fundamental cellular processes such as transcription, replication, and DNA repair (1). The basic structural unit of chromatin is the nucleosome, which comprises approximately 147 base pairs of DNA wrapped around a histone octamer containing two copies each of histones H2A, H2B, H3, and H4 (2, 3). The positions in nucleosomal DNA are designated as superhelical locations (SHLs) (4). By defining the center of the nucleosomal DNA as SHL(0), which corresponds to the dyad axis of the nucleosome, DNA positions at roughly every 10 base pairs are assigned as SHL(±1) to SHL(±7) (2, 4). The dynamic nature of nucleosomes is tightly regulated by post-translational modifications (PTMs) of histones, which can modulate histone-DNA and histone-protein interactions to influence chromatin accessibility (5–7). Among these modifications, the acetylation of lysine residues on histone H3 has long been associated with transcriptional activation, by promoting an open chromatin conformation conducive to gene expression (8, 9).

While histone acetylation is generally thought to weaken histone-DNA contacts by neutralizing the positive charge of lysine side chains (10), how individual acetylation sites contribute to chromatin dynamics and transcriptional regulation remains poorly understood. In particular, the lysine residues within the histone H3 core domain, such as Lys56 (K56), Lys64 (K64), and Lys122 (K122), are positioned at key DNA-histone interfaces and have been implicated in modulating nucleosome stability (6, 11–13). Mutations at these residues have been shown to reduce viability and fertility in *Drosophila* (14, 15). H3K56 is located at the DNA entry/exit region and is acetylated by p300/CBP and deacetylated by SIRT1, SIRT2, and SIRT6 (16–19). Rtt109, a fungal histone acetyltransferase with a p300/CBP-like catalytic fold, acetylates H3K56 in H3-H4 complexes, whereas no detectable activity toward nucleosomal H3 has been observed, suggesting that H3K56ac is introduced prior to or during nucleosome deposition (20). H3K56ac has been linked to nucleosome assembly, disassembly, and replication-coupled chromatin remodeling (21–24). In addition, H3K56ac contributes to transcriptional regulation. H3K56ac is enriched in transcription start sites (TSS) of highly expressed genes (25), and a yeast strain carrying the acetylation-mimic mutation H3K56Q exhibits increased survival in the presence of the transcription inhibitor 6-azauracil (26). The acetylation of H3K122, located near the nucleosomal dyad, is mediated by p300/CBP (27). This acetylation has been reported to directly destabilize histone-DNA contacts, promoting nucleosome sliding and transcriptional activation (27–31). H3K122ac is enriched at TSS and enhancers of active genes (27, 32). Similarly, the acetylation of H3K64, a residue located within the nucleosome core, is also enriched at TSS and enhancers of active genes and is catalyzed by p300/CBP (13). This modification destabilizes nucleosomes and enhances chromatin accessibility (13, 32). However, the mechanisms by which these modifications alter the nucleosome architecture to facilitate RNA polymerase II (RNAPII) transcription remain elusive.

In the present study, we found that the acetylation of histone H3 at Lys56 and Lys122 markedly enhances RNA polymerase II (RNAPII) transcription through nucleosomes, whereas the acetylation at Lys64 has little effect. To elucidate the structural basis for these functional differences, we determined cryo-electron microscopy (cryo-EM) structures of nucleosomes containing site-specifically acetylated H3 at Lys56, Lys64, or Lys122. The structures revealed that H3K56ac and H3K122ac locally weaken histone-DNA interactions at the DNA entry/exit region and near the dyad, respectively, whereas H3K64ac causes no detectable changes. These distinct structural consequences of acetylation within the histone fold domain of H3 provide a mechanistic framework for understanding how specific lysine modifications fine-tune nucleosome dynamics to facilitate RNAPII transcription.

## Results

### Preparation of site-specifically acetylated nucleosomes for RNAPII transcription

To investigate how specific acetylation sites within the histone fold domain of histone H3 influence nucleosome structure and transcription, we prepared human histone H3 bearing the site-specific acetylation of Lys56, Lys64, or Lys122 (H3K56ac, H3K64ac, and H3K122ac) by chemical synthesis (Figure 1A and B, Supplementary Figure 1A and B). Each acetylated histone H3 was obtained through native chemical ligation with synthetic acetylated and non-acetylated peptides. In this method, we employed the H3.2 C110A mutant, in which the Cys110 residue of H3.2 was replaced by Ala, because the Cys residue is converted into the Ala residue during the desulfurization process after peptide ligation. Histone H3.1 contains an additional cysteine at residue 96, whereas the corresponding residue in H3.2 is serine. Therefore, H3.2 was selected to avoid unintended conversion of Cys96 to alanine during desulfurization and thereby minimize additional sequence alterations. We prepared histone octamers containing recombinant histones H2A, H2B, H4, and the synthesized H3.2 containing H3K56ac, H3K64ac, or H3K122ac. Nucleosomes containing H3K56ac, H3K64ac, and H3K122ac were then reconstituted using a 198 base-pair (bp) DNA fragment consisting of a modified Widom 601 sequence containing a 9-base mismatched region in a linker DNA (33) (Figure 1B, Supplementary Figure 1C and D). RNAPII associates with the 9-base mismatched region and initiates transcription by the addition of nucleotide triphosphates and a primer RNA. We then performed in vitro nucleosome transcription assays using purified *Komagataella phaffii* RNAPII and its essential elongation factor, TFIIS (Figure 1C).

**Figure 1:**
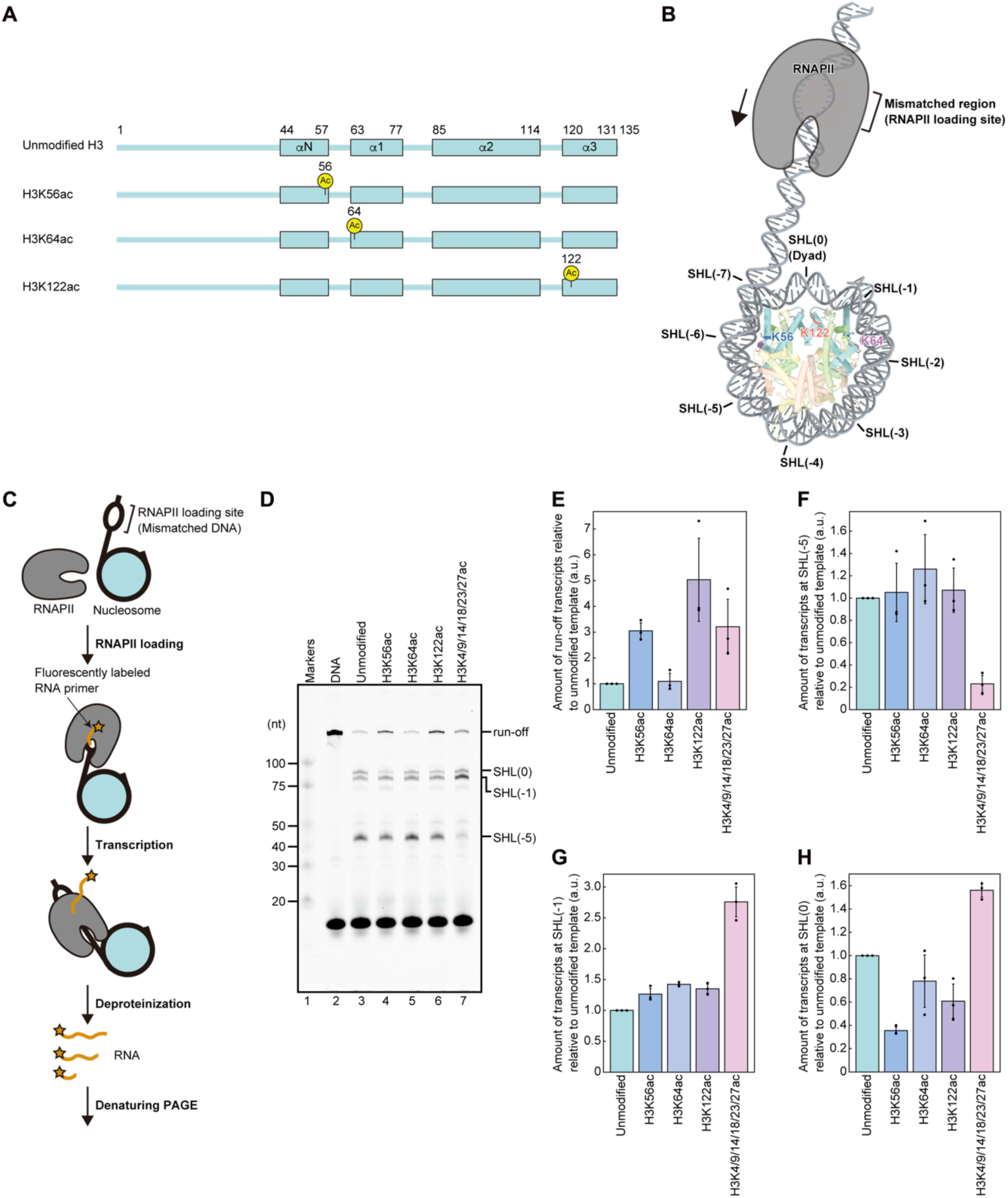
Acetylation at Lys56 or Lys122 of H3 decreases transcription barrier at SHL(0) and facilitates RNAPII progression. (A) Secondary structure of histone H3, showing the acetylated residues (yellow). (B) SHLs, transcription direction, and Lys56, Lys64, and Lys122 are shown on the nucleosome structure (modified from PDB ID: 7K61). The linker DNA contains a mismatched region that serves as the RNAPII loading site. Histones H2A, H2B, H3, H4, and DNA are colored pale yellow, pale red, pale blue, pale green, and gray, respectively. Lys56, Lys64, and Lys122 are shown in blue, purple, and orange, respectively. (C) Schematic representation of the transcription assay. (D) Transcription assays using free DNA, and unmodified, H3K56ac, H3K64ac, H3K122ac, and H3K4/9/14/18/23/27ac nucleosome templates. RNA transcripts were analyzed by denaturing PAGE and detected by DY647 fluorescence. Transcription reactions were terminated after 30 min. (E-H) Quantification of transcripts corresponding to (E) run-off, (F) SHL(−5), (G) SHL(−1), and (H) SHL(0) positions. Amounts of transcripts were normalized to those of the unmodified nucleosome template. Bar graphs show the means of three independent experiments, and dots indicate the individual values. Error bars represent the standard deviation. No statistical testing or multiple-comparison correction was applied.

### Site-specific acetylations of the histone fold domain of H3 differently modulate nucleosome transcription by RNAPII

When RNAPII transcribed through the unmodified nucleosome, an RNA transcript of approximately 40 nucleotides, which corresponds to the RNAPII pausing at the SHL(−5), was predominantly accumulated (Figure 1D, lane 3, Supplementary Figure 2). In addition, RNA products around 75-80 nucleotides, which correspond to the RNAPII pausing at the SHL(−1) and SHL(0) positions, were also detected (Figure 1D, lane 3). Notably, run-off transcripts were barely detectable in the unmodified nucleosome template (Figure 1D, lane 3). These results are consistent with the previous studies (33) and indicate that the nucleosome acts as a substantial barrier to RNAPII progression.

Strikingly, nucleosomes containing either H3K56ac or H3K122ac showed a pronounced increase in run-off transcript production by RNAPII (Figure 1D, lanes 4, 6). In contrast, H3K64ac had little or no effect on the transcription efficiency, which was comparable to the unmodified control (Figure 1D, lane 5). These results demonstrate that specific acetylations within the histone fold domain distinctly modulate the transcription efficiency of RNAPII through the nucleosome.

A quantitative analysis revealed that H3K56ac and H3K122ac increased the fractions of full-length transcripts by approximately three- and five-fold relative to unmodified nucleosomes, respectively (Figure 1E). These effects are comparable to the enhanced nucleosome transcription observed with the H3 N-terminal acetylations, including H3K4ac, H3K9ac, H3K14ac, H3K18ac, H3K23ac, and H3K27ac (Figure 1D, lane 7, and E). Consistent with a previous study (34), the H3 N-terminal acetylations drastically reduce RNAPII pausing at the SHL(−5) position, thereby increasing the pausing population at the SHL(−1) and SHL(0) positions (Figure 1F-H). In contrast, the RNAPII pausing at the SHL(−5) and SHL(−1) positions was not substantially altered in the H3K56ac and H3K122ac nucleosomes (Figure 1F and G). Notably, the RNAPII pausing at the SHL(0) position was significantly alleviated in the H3K56ac and H3K122ac nucleosomes (Figure 1H). Such alleviation of RNAPII pausing at the SHL(0) position was not observed in the nucleosome containing the H3 N-terminal acetylations (Figure 1D, lane 7, and H). Therefore, the acetylation within the histone fold domain may regulate RNAPII transcription differently from the acetylations on the H3 N-terminal tail.

### Cryo-EM structures reveal local weakening of histone-DNA contacts by H3K56ac and H3K122ac, but not H3K64ac

To understand the structural basis for the distinct transcriptional outcomes, we determined the cryo-EM structures of the H3K56ac, H3K64ac, and H3K122ac nucleosomes and the unmodified nucleosome used for the nucleosome transcription assays at 2.96, 2.99, 3.07, and 3.07 Å resolutions, respectively (Figure 2A, Table 1, and Supplementary Figures 3-7). We avoided sample crosslinking, which stabilizes histone-DNA contacts in the nucleosome. Instead, we employed a single-chain variable fragment (scFv), PL2-6, which specifically binds to the acidic patch of nucleosomes without contacting the nucleosomal DNA and effectively isolates the nucleosome sample from the air-water interface, thereby stabilizing the cryo-EM specimen (35).

**Figure 2:**
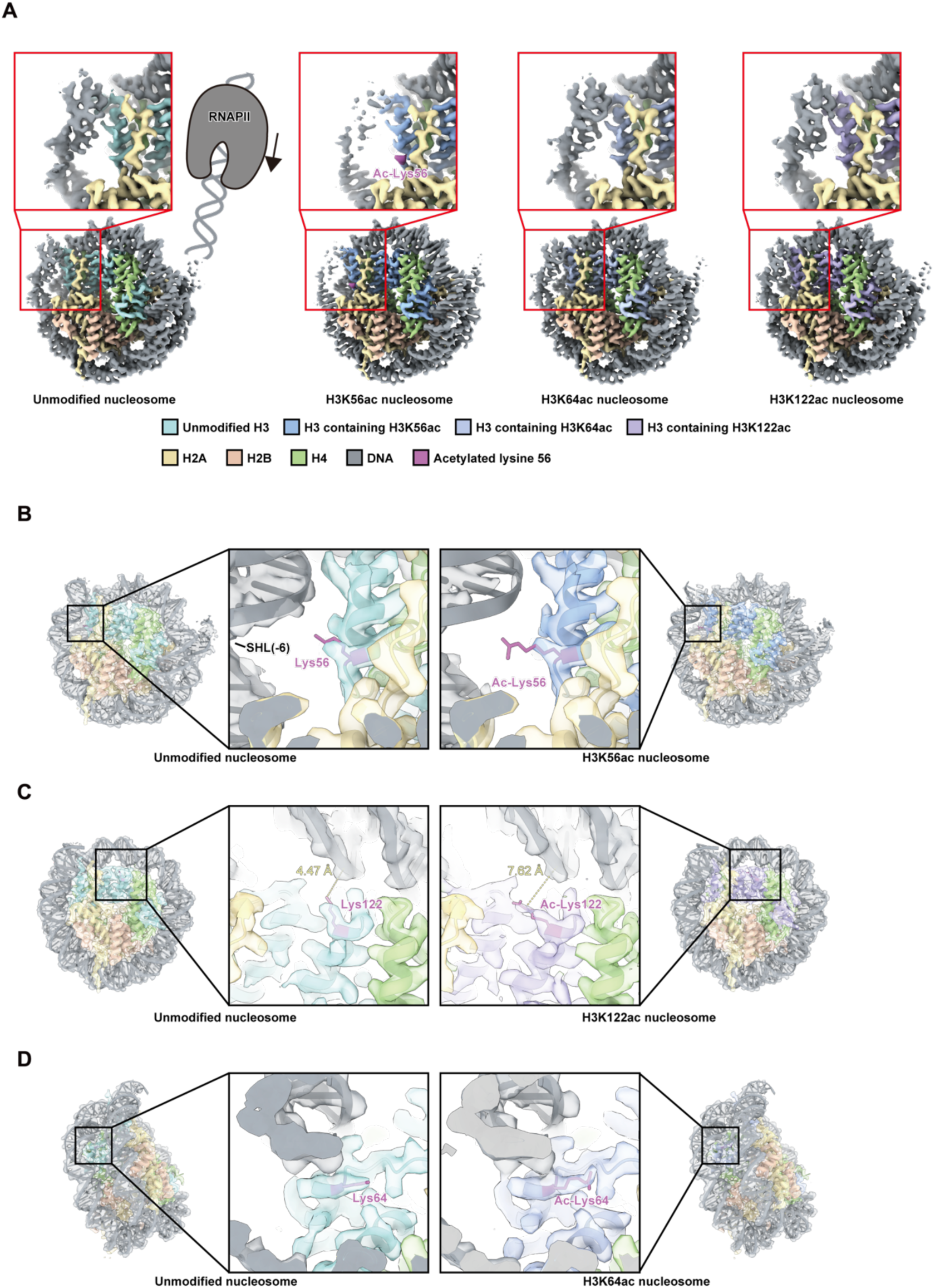
Cryo-EM structures reveal weakened histone–DNA interactions induced by H3K56ac and H3K122ac. (A) Cryo-EM density maps of the unmodified, H3K56ac, H3K64ac, and H3K122ac nucleosomes. H2A, H2B, H4, and DNA are colored yellow, red, green, and gray, respectively. H3 in the unmodified, H3K56ac, H3K64ac, and H3K122ac nucleosomes is colored cyan, blue, dark blue, and purple, respectively. The density map corresponding to Ac-Lys56 is shown in magenta. Cryo-EM maps were low-pass filtered to 3.1 Å and normalized by root-mean-square scaling relative to the unmodified nucleosome map. All maps are displayed at a contour level of 0.013. (B-D) Close-up views of the regions surrounding the acetylated residues in H3K56ac (B), H3K122ac (C), and H3K64ac (D) nucleosomes. H2A, H2B, H4, and DNA are colored yellow, red, green, and gray, respectively. H3 in the unmodified, H3K56ac, H3K64ac, and H3K122ac nucleosomes is colored cyan, blue, dark blue, and purple, respectively. Acetylated lysine residues and their corresponding unmodified lysine residues are shown in magenta. Cryo-EM maps were low-pass filtered to 3.1 Å and normalized by root-mean-square scaling relative to the unmodified nucleosome map. All maps are displayed at a contour level of 0.01. The distances between the nitrogen atom at the zeta position of the lysine or acetyl-lysine residue in chain E and the nearest oxygen atom of the DNA backbone in chain J are shown in (C).

**Table 1.** Cryo-EM data collection, processing, refinement, and validation statistics.

| Sample | #1 Structure of the unmodified nucleosome (EMD-66767) (PDB ID: 9XDM) | #2 Structure of the H3K56ac nucleosome (EMD-66768) (PDB ID: 9XDN) | #3 Structure of the H3K64ac nucleosome (EMD-66769) (PDB ID: 9XDO) | #4 Structure of the H3K122ac nucleosome (EMD-66770) (PDB ID: 9XDP) |
|---|---|---|---|---|
| **Data collection** |  |  |  |  |
| Electron microscope | KriosG4 | KriosG4 | KriosG4 | KriosG4 |
| Camera | K3 | K3 | K3 | K3 |
| Pixel size (Å/pix) | 1.06 | 1.06 | 1.06 | 1.06 |
| Defocus range (µm) | -1.25 to -2.25 | -1.25 to -2.25 | -1.25 to -2.25 | -1.25 to -2.25 |
| Exposure time (second) | 4.5 | 4.5 | 4.5 | 4.5 |
| Total dose (e<sup>-</sup>/Å<sup>2</sup>) | 61.9 | 60.9 | 60.6 | 61.4 |
| Movie frames (no.) | 40 | 40 | 40 | 40 |
| Total micrographs (no.) | 6 580 | 8 180 | 8 071 | 7 431 |
| **Reconstruction** |  |  |  |  |
| Software | Relion 5.0 | Relion 5.0 | Relion 5.0 | Relion 5.0 |
| Particles for 2D classification | 2 832 919 | 4 165 413 | 2 752 619 | 4 247 723 |
| Particles for 3D classification | 684 801 | 811 976 | 561 678 | 669 653 |
| Particles in the final map (no.) | 91 479 | 122 391 | 124 333 | 127 356 |
| Symmetry | C1 | C1 | C1 | C1 |
| Final resolution (Å) | 3.07 | 2.96 | 2.99 | 3.07 |
| FSC threshold | 0.143 | 0.143 | 0.143 | 0.143 |
| Map sharpening B factor (Å<sup>2</sup>) | -71.248 | -71.827 | -83.214 | -82.055 |
| **Model building** |  |  |  |  |
| Software | Coot | Coot | Coot | Coot |
| **Refinement** |  |  |  |  |
| Software | Phenix, ISOLDE | Phenix, ISOLDE | Phenix, ISOLDE | Phenix, ISOLDE |
| **Model composition** |  |  |  |  |
| Protein | 726 | 726 | 726 | 721 |
| Nucleotide | 268 | 262 | 266 | 264 |
| **Validation** |  |  |  |  |
| MolProbity score | 1.28 | 0.96 | 1.28 | 1.27 |
| Clash score | 5.23 | 1.93 | 5.25 | 5.03 |
| R.m.s. deviations |  |  |  |  |
| Bond lengths (Å) | 0.004 | 0.005 | 0.004 | 0.004 |
| Bond angles (˚) | 0.759 | 0.674 | 0.761 | 0.784 |
| Ramachandran plot Z-score |  |  |  |  |
| Whole | 0.79 | 1.03 | 0.79 | 1.06 |
| Helix | 0.95 | 1.20 | 0.97 | 1.16 |
| Sheet | n/a | n/a | n/a | n/a |
| Loop | -0.51 | -0.63 | -0.57 | -0.49 |
| CaBLAM outliers (%) | 1.15 | 1.01 | 1.15 | 1.02 |
| **Ramachandran plot** |  |  |  |  |
| Favored (%) | 98.31 | 98.16 | 98.30 | 98.28 |
| Allowed (%) | 1.55 | 1.84 | 1.56 | 1.57 |
| Outliers (%) | 0.14 | 0.00 | 0.14 | 0.14 |
| **Map to model correlation** |  |  |  |  |
| Masked correlation coefficient | 0.86 | 0.86 | 0.84 | 0.82 |

In the H3K56ac nucleosome, parts of the Lys56 and acetylated Lys56 side chains are resolved, with the visible density appearing to extend toward the SHL(±6) region (Figure 2B). In addition, the density corresponding to the DNA entry/exit region around SHL(±6) appeared substantially weaker than that of the unmodified nucleosome, suggesting the increased flexibility of the nucleosomal DNA ends. In contrast, the density differences at SHL(±5) and SHL(0) were not as pronounced (Figure 2A, Supplementary Figure 8). Since lysine acetylation diminishes its positive charge and thereby weakens the local histone-DNA interactions, the flexibility of the nucleosomal DNA ends may be enhanced by the H3 Lys56 acetylation, which weakens the histone-DNA interactions near the SHL(±6) position despite the overall preservation of the side-chain orientation. Because cryo-EM analysis may preferentially retain more ordered particles, we also performed biochemical assays to assess the flexibility of the nucleosomal DNA ends. Consistently, the DNA ends of the H3K56ac nucleosome were markedly more susceptible to micrococcal nuclease than those of the unmodified nucleosome (Figure 3A-C, Supplementary Figure 9). These results are consistent with previous fluorescence resonance energy transfer studies (29, 36–38). Notably, the leading edge of RNAPII sterically clashes with the distal DNA end of the nucleosome upon reaching the SHL(0) position, inducing RNAPII pausing (39). Therefore, the increased flexibility at the distal nucleosomal DNA end may alleviate the RNAPII pausing at the SHL(0) position (Figure 4).

**Figure 3:**
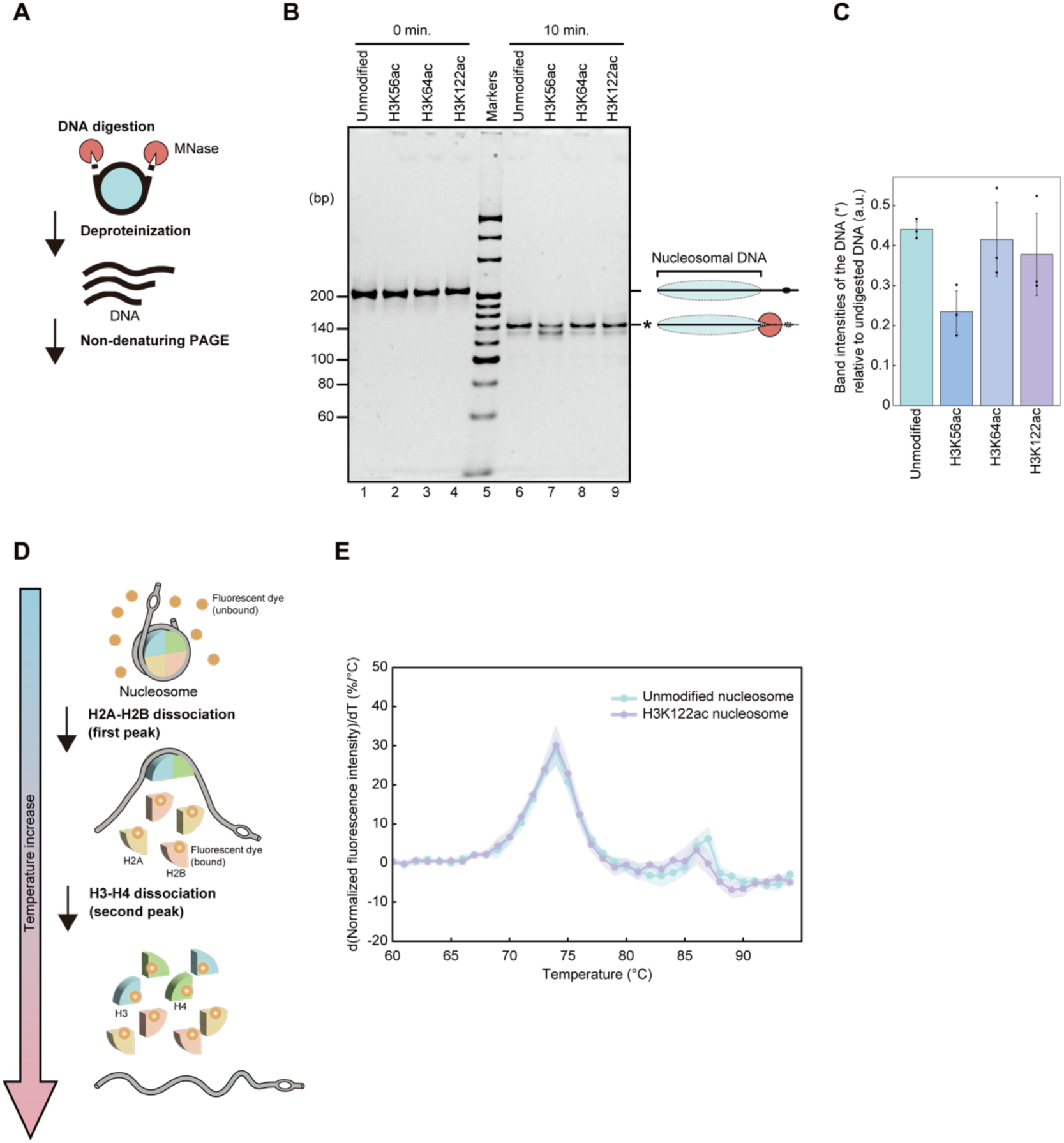
Biochemical analyses support weakened histone-DNA interactions induced by H3K56ac and H3K122ac. (A) Schematic representation of the micrococcal nuclease digestion assay. (B) Micrococcal nuclease digestion assays of the unmodified, H3K56ac, H3K64ac, and H3K122ac nucleosomes. Deproteinized DNA was analyzed by non-denaturing PAGE and stained with ethidium bromide. (C) Quantification of the micrococcal nuclease assay. Band intensities indicated by asterisks in (B) were normalized with those of the undigested DNA. Bar graphs show the means of three independent experiments, and dots indicate the individual values. Error bars represent the standard deviation. No statistical testing or multiple-comparison correction was applied. (D) Schematic representation of the thermal stability assay. (E) Thermal stability assay of unmodified and H3K122ac nucleosomes. The line graph shows the mean values from three independent experiments, and the shaded areas represent the standard deviations (SD). No statistical testing or multiple-comparison correction was applied.

**Figure 4:**
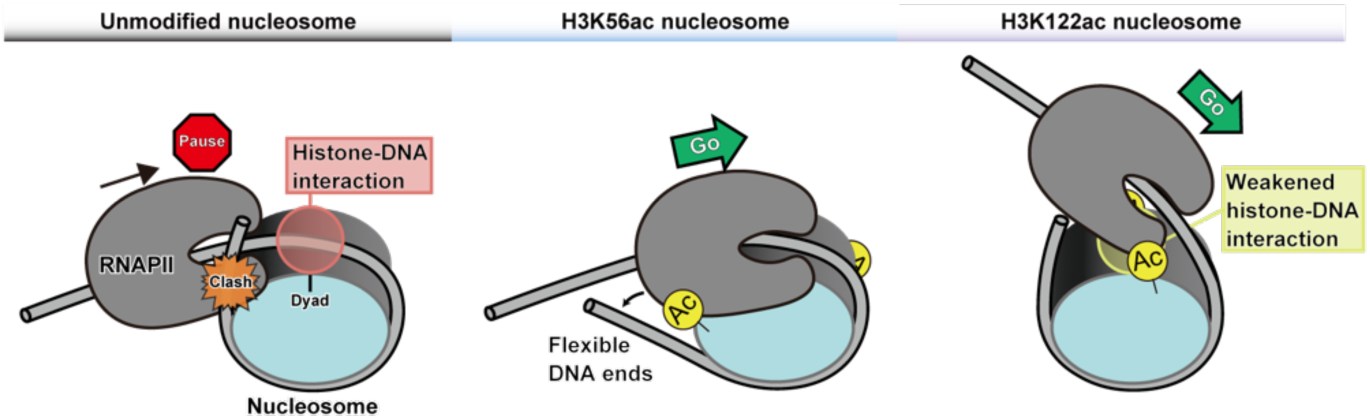
Acetylation at Lys56 and Lys122 of H3 likely promotes transcription through distinct yet complementary mechanisms. The direct contacts between the distal DNA end of the nucleosome and RNAPII, as well as the histone-DNA interactions near the dyad axis, likely contribute to RNAPII pausing at the SHL(0) position. The acetylation at Lys56 of H3 enhances the flexibility of the DNA ends of the nucleosome, which may alleviate steric clashes between the DNA and RNAPII and facilitate RNAPII passage. In contrast, the acetylation at Lys122 of H3 weakens histone-DNA interactions near the dyad axis, thereby reducing RNAPII pausing and enhancing transcription at the SHL(0) region.

The H3 Lys122 residue directly interacts with the DNA backbone in the unmodified nucleosome. The acetyl group neutralizes the positive charge of the lysine sidechain, thereby diminishing the electrostatic attraction between the histone and DNA, which likely facilitates the partial unwrapping of the nucleosomal DNA near the SHL(0) position. The structural comparison suggests that acetylation of the H3 Lys122 residue reorients the side chain away from the nucleosomal DNA, with this change being most evident in one of the two H3 copies within the nucleosome (Figure 2C). To assess the effect of H3K122ac on nucleosome stability, we performed a thermal stability assay (Figure 3D and E, Supplementary Figure 10). In this assay, nucleosomes were mixed with a fluorescent dye whose fluorescence intensity increases upon binding to hydrophobic regions exposed during thermal denaturation. The first transition corresponding to H2A-H2B dissociation was similar between the unmodified and H3K122ac nucleosomes, whereas the second transition corresponding to H3-H4 dissociation was approximately 1°C lower in the H3K122ac nucleosome (Figure 3E). This is consistent with the previously reported thermal stability assay of the nucleosome-like complex of H3-H4-DNA bearing H3K122ac (31). These structural and biochemical observations support a model in which H3K122ac weakens histone-DNA interactions around SHL(0), thereby enhancing RNAPII transcription by alleviating pausing at SHL(0) (Figure 4).

In contrast, the H3K64ac nucleosome displayed nearly identical density and local resolution profiles to the unmodified nucleosome, indicating that this modification does not significantly alter the histone-DNA contacts within the nucleosome core particle (Figure 2D).

## Discussion

Our results reveal that the acetylation of lysine within the histone fold domain of H3 has distinct and site-specific effects on RNAPII transcription through nucleosomes. Among the three tested sites, H3K56ac and H3K122ac markedly enhanced transcription, whereas H3K64ac had little influence. Structural analyses by cryo-EM demonstrated that H3K56ac and H3K122ac locally weaken the histone-DNA interactions at the DNA entry/exit region and near the dyad, respectively, consistent with their effects on transcription. These findings establish a mechanistic link between the site-specific acetylation within the histone fold domain and the regulation of RNAPII transcription by histone acetylation.

The acetylations of the H3 Lys56 and H3 Lys122 residues likely promote transcription by distinct but complementary mechanisms (Figure 4). H3K56ac increases the flexibility of nucleosomal DNA ends, which may lower the energetic barrier for DNA unwrapping and alleviate the RNAPII pausing at the SHL(0) position by reducing steric clashes between RNAPII and the distal nucleosomal DNA end. In contrast, H3K122ac destabilizes the histone-DNA contacts around the dyad, directly weakening the central anchor point of the nucleosome and facilitating the passage of RNAPII through this region. Both effects converge to lower the nucleosomal barrier and enhance transcriptional output, consistent with the observed increases in full-length transcripts. Notably, these mechanisms differ from those mediated by H3 N-terminal acetylations, which mainly affect RNAPII pausing at upstream positions such as SHL(−5) (34). Thus, the acetylation within the histone fold domain modulates transcription through a structural mechanism distinct from that of histone tail acetylations. Previous biochemical studies have suggested that H3K56ac and H3K122ac contribute to chromatin remodeling, histone exchange, and transcriptional activation (27, 28, 30, 40, 41). Our structural data now provide direct evidence that these modifications alter the physical properties of the nucleosome to promote transcriptional elongation. Future structural analyses of RNAPII complexes with acetylated nucleosomes may provide further mechanistic insights into this process.

Although H3K64ac is enriched at transcriptionally active sites, our results indicate that acetylation of H3 Lys64 has minimal impact on RNAPII transcription through nucleosomes or on nucleosome structure. Our transcription system includes RNAPII and TFIIS but no additional transcription-associated factors, allowing us to isolate the direct effects of individual histone acetylation events. Di Cerbo et al. have reported that H3K64ac promotes histone eviction, raising the possibility that additional factors, such as histone chaperones, are required to facilitate transcription (13). This difference raises the possibility that H3K64ac influences transcription through histone exchange or other chromatin remodeling processes involving additional factors, such as histone chaperones. Moreover, H3K64ac is known to co-localize with transcriptionally active histone marks, including H3K27ac, H3K4me1, and H3K122ac, suggesting that H3K64ac may regulate transcription through crosstalk with other PTMs (13, 32). In our H3K64ac nucleosome structure, acetylated Lys64 is shielded by the surrounding DNA, implying that this modification is inaccessible to reader proteins in an intact nucleosome. Therefore, recognition of H3K64ac likely requires nucleosome disruption. Crosstalk between H3K64ac and other PTMs may occur prior to chromatin assembly or during nucleosome-disruptive events. The roles of additional factors and PTM crosstalk remain important subjects for future investigation.

More broadly, our findings highlight the functional diversity of histone acetylation beyond the N-terminal tails. Whereas the N-terminal tail acetylations predominantly recruit effector proteins through bromodomains and other reader modules, histone fold domain acetylations act directly on nucleosomal DNA dynamics (6, 8, 42). This dual mode of regulation, via effector-mediated signaling and direct modulation of nucleosome architecture, may enable cells to precisely coordinate chromatin remodeling and transcription in response to specific cues (43). Our work provides a foundation for future studies to investigate how distinct histone fold domain acetylations interact with other chromatin modifications, remodeling factors, and transcription machinery across different physiological and developmental contexts, thereby advancing our understanding of how the nucleosome structure encodes regulatory information.

## Material and methods

### Preparation of histones

Human histones H2A, H2B, and H4 were expressed and purified as described previously (44). Briefly, H2A, H2B, and H4 were expressed in Escherichia coli cells. After denaturation of the insoluble fractions, the 6×histidine-tagged histones were purified by Ni-NTA agarose chromatography (Qiagen). The 6×histidine-tags were removed by thrombin protease treatment and the histones were further purified by MonoS cation-exchange chromatography.

Histone H3.2 C110A was used as the common background for the unmodified, H3K56ac, H3K64ac, and H3K122ac histones prepared by native chemical ligation. The C110A substitution was introduced to eliminate the only native cysteine residue in H3.2 and thereby prevent undesired side reactions during the desulfurization step.

Unmodified human histone H3.2 C110A was purified as previously described (45). In summary, the chemically synthesized H3 N-terminal peptide (residues 1-28) was ligated to the recombinant C-terminal regions of H3 (A29C-135, C110A) using the one-pot native chemical ligation method. After desulfurization, the H3 peptide was purified by high performance liquid chromatography. Because cysteine at position 29 was converted to alanine during desulfurization, the resulting peptide contained only the C110A mutation.

The human histone H3.2 C110A acetylated at Lys 56 (H3K56ac) was previously synthesized (46) and used in this study. The other acetylated histones (H3K64ac and H3K122ac) were newly synthesized in this study by total chemical synthesis, as described below.

The full-length H3.2 sequence was divided into three peptide segments (i.e., N-terminal, internal, and C-terminal segments). Each segment was prepared through Fmoc solid-phase peptide synthesis (SPPS): the N-terminal segment as an N-terminal free amine with a C-terminal thioester, the internal segment as an N-terminal thiazolidine (Thz) with a C-terminal thioester, and the C-terminal segment as an N-terminal Cys with a C-terminal carboxylic acid. Automated SPPS was carried out on a MultiPep RS synthesizer (Intavis) according to the standard Fmoc-SPPS protocol. Coupling was performed twice with Fmoc-protected amino acids (5.2 equiv), HBTU (5.0 equiv), and DIEA (10.0 equiv) in DMF for at least 20 min each at room temperature, and Fmoc removal was performed twice with 20% (v/v) piperidine in DMF for 5 min each at room temperature. The peptide thioester segments (i.e., N-terminal and internal segments) were synthesized on 2-chlorotrityl chloride resin, onto which Fmoc-NHNH₂ was loaded prior to chain elongation, whereas the peptide acid segment (i.e., C-terminal segment) was synthesized on Fmoc-Ala-Wang-PEG resin. For acetylated Lys incorporation, Fmoc-Lys(Ac)-OH was coupled at position 64 in the internal segment or position 122 in the C-terminal segment. Following chain elongation, cleavage from the resin and global deprotection were performed for 2 h using TFA/TIS/H₂O (95:2.5:2.5) for the N-terminal segment, and TFA/EDT/anisole/thioanisole (90:3:2:5) supplemented with 10% (v/v) dimethyl sulfide and 1% (w/v) TBAI for the internal and C-terminal segments. The N-terminal and internal segments were converted into the corresponding C-terminal thioesters prior to HPLC purification. For the N-terminal segment, the crude peptide hydrazide was dissolved in 8 M Gn·HCl buffer (pH 3.0), and the solution was cooled to −15°C. NaNO₂ was then added to oxidize the hydrazide to the corresponding acyl azide. MESNa was subsequently added, and the pH was adjusted to 6.0, giving the peptide MESNa thioester. For the internal segment, the crude peptide hydrazide was dissolved in 8 M Gn·HCl buffer (pH 3.0), and acetylacetone (2.5 equiv) and MPAA (final 100 mM) were added. The mixture was incubated at 37°C for 1 h to afford the peptide MPAA thioester. After purification of crude peptides by preparative HPLC, each peptide was checked by analytical HPLC for purity, and was identified by MALDI-TOF MS.

For peptide ligation, the internal segment (1.1 equiv.) and the C-terminal segment (1.0 equiv.) were dissolved at 1.5 mM and 1.4 mM, respectively, in buffer containing 8 M guanidine hydrochloride (Gn·HCl), 100 mM 4-mercaptophenylacetic acid (MPAA), 20 mM TCEP·HCl and 0.2 M NaH₂PO₄ (pH 7.0). The mixture was stirred at 37°C for 2.5 h under an argon atmosphere. MeONH2·HCl was then added to a final concentration of 100 mM, and the pH of the reaction mixture was adjusted to 4.0 by adding 6 M HCl to convert the N-terminal Thz into cysteine. After stirring for 23 h, the N-terminal segment (1.2 equiv.) and additional TCEP·HCl (20 mM, pH 7.0) were added, and the pH of the solution was readjusted to 7.0 with 1 M NaOH. The resulting mixture was stirred at 37°C for 15 h under an argon atmosphere, with the reaction progress monitored by analytical HPLC and MALDI-TOF MS. The ligated product was purified by preparative HPLC and identified by MALDI-TOF MS.

For desulfurization, the ligated peptide was dissolved in degassed buffer containing 8 M guanidine hydrochloride (Gn·HCl) and 0.2 M NaH₂PO₄ (pH 7.0). Then, TCEP·HCl, glutathione and VA-044 were added to give final concentrations of 0.5 mM peptide, 300 mM TCEP, 80 mM glutathione, and 20 mM VA-044. The pH of the reaction mixture was adjusted to 7.0, and the mixture was stirred at 37°C for 9 hours under an argon atmosphere. After purification of the desulfurized products by preparative HPLC, the isolated products were checked by SDS-PAGE for purity, and identified by MALDI-TOF MS.

### Preparation of DNA fragment

The 198 bp DNA fragment containing the 9-base mismatched region was purified as previously described (33). In brief, the plasmid containing the modified Widom 601 DNA fragment was amplified in *E. coli*. The Widom 601 DNA fragment was excised from the plasmid DNA by *Eco*RV (Takara), and purified by polyethylene glycol precipitation. After dephosphorylation with alkaline phosphatase (Takara), the DNA fragment was digested with *Bgl*I (Takara). The resulting fragment was purified by a Prep Cell apparatus (Bio-Rad). The purified 153 bp DNA fragment was ligated with the 45 bp DNA fragment containing the 9-base mismatched region by T4 DNA ligase (NIPPON GENE), and further purified by a Prep Cell apparatus (Bio-Rad).

### Reconstitution of nucleosomes

The nucleosomes were reconstituted as previously reported (44). In brief, the histone octamers were assembled from H2A, H2B, H4, and either unmodified or acetylated H3 histones, and purified by size-exclusion chromatography on a HiLoad 16/60 Superdex 200 column (Cytiva). The resulting octamers were mixed with the 198 bp DNA fragment, and the nucleosomes were reconstituted using the salt-dialysis method. The nucleosomes were further purified by a Prep Cell apparatus (Bio-Rad).

### Preparation of RNAPII and TFIIS

*Komagataella phaffii* RNAPII was purified as previously described (47–49). In summary, endogenous Rpb2 containing the tandem affinity purification (TAP) tag was purified from *K. phaffii* cells by FLAG-affinity column chromatography and anion-exchange column chromatography.

*K. phaffii* TFIIS was prepared as previously described (49). Briefly, the TFIIS protein was bacterially produced and purified by Ni-affinity column chromatography. After removing the His-tag by HRV-3C protease, the TFIIS protein was further purified by cation-exchange column chromatography.

### Transcription assay

The transcription assay was conducted as previously described (33, 34). In brief, the nucleosome and RNAPII were incubated at 30°C for 30 minutes in reaction buffer, containing 0.1 µM nucleosome, 0.1 µM RNAPII, 0.1 µM TFIIS, 26 mM HEPES-KOH (pH 7.5), 50 mM potassium acetate, 0.2 µM zinc acetate, 20 µM Tris(2-carboxyethyl) phosphine, 1.5% glycerol, 0.1 mM dithiothreitol, 5 mM magnesium chloride, 400 µM each ATP, UTP, GTP, and CTP, and 0.4 µM fluorescently labeled RNA primer (5′-DY647-AUAAUUAGCUC-3′) (Dharmacon). The reaction was quenched by mixing 2 µl of the reaction mixture and 1 µl of deproteinization solution containing 200 mM Tris-HCl (pH 8.0), 80 mM EDTA, and 0.6 mg/ml proteinase K (Roche), followed by an incubation at room temperature for 10 minutes. Then, 12 µl of Hi-Di formamide was added, and the sample was heated at 95°C for 5 minutes. The sample was analyzed by 10% denaturing PAGE. The DY647 fluorescence of RNA products was detected by an Amersham Typhoon imager (Cytiva). The band intensities of the transcripts were quantified with the ImageQuant™ TL software (GE Healthcare) and normalized with the transcripts of the unmodified nucleosome template. The transcription assay was performed in triplicate, and the mean values are presented with standard deviations (SD) shown as error bars. No statistical testing or multiple-comparison correction was applied.

### Purification of PL2-6 scFv

PL2-6 scFv containing (GGGS)4 linker peptides between the heavy and light chains was purified as previously described (34, 35, 50). In short, the scFv was bacterially produced and recovered as an insoluble fraction. After denaturation, the histidine-tagged scFv was purified by Ni-NTA agarose chromatography (Qiagen). The resulting scFv was refolded by dialysis with buffer exchange using a peristaltic pump, and further purified on a HiLoad 26/600 Superdex 75 pg column (Cytiva). Eluted fractions were analyzed by SDS-PAGE under both reducing conditions with an incubation at 95°C for 5 minutes and non-reducing conditions without heating. The binding activity of the scFv in each fraction was assessed by an electrophoretic mobility shift assay using nucleosomes. Fractions exhibiting nucleosome-binding activity were combined, concentrated, and stored at −80°C.

### Sample preparation for cryo-EM analysis

Samples of unmodified H3, H3K56ac, H3K64ac, and H3K122ac nucleosomes for cryo-EM analysis were prepared as previously described (34). Nucleosomes (0.5 µM) and 1.5 µM of PL2-6 scFv were mixed in reaction buffer, containing 10 mM HEPES-KOH (pH 7.5), 0.5 mM dithiothreitol, 2.5% glycerol, 10 mM Tris-HCl (pH 7.5), 30 mM sodium chloride, and 0.2 mM EDTA. After an incubation at 30°C for 30 minutes, the mixture was applied to a Quantifoil R1.2/1.3 copper grid and plunge-frozen using a Vitrobot Mark IV (Thermo Fisher Scientific). Grids were glow discharged for 1 minute using a PIB-10 Bombarder (Vacuum Device Inc.) prior to sample application.

### Cryo-EM data collection

The cryo-EM data were acquired using the EPU automation software on a Krios G4 electron microscope (Thermo Fisher Scientific). The microscope was operated at 300 kV, with a nominal magnification of 81,000× (corresponding to a pixel size of 1.06 Å), and a defocus range of −1.25 to −2.25 µm. Micrographs were recorded on a K3 BioQuantum direct electron detector (Gatan) with a 20 eV energy-filter, with a total of 40 frames per movie. The detailed cryo-EM data recording conditions are provided in Table 1.

### Image processing

The overall workflow is shown in Supplementary Figure 3-6. The acquired movies were aligned and dose-weighted using MOTIONCOR2 (51). CTFFIND4 (52) was used to estimate the contrast transfer function (CTF). Relion 5.0 (53) was used for image processing. Particles were picked by Laplacian-of-Gaussian based auto picking and subjected to two-dimensional classification, de novo initial model generation, three-dimensional (3D) classification, 3D auto-refinement, Bayesian polishing, and CTF refinement. Junk particles were removed during the classification steps. To focus on the nucleosome structure, 3D classification with a nucleosome mask was conducted with and without alignment. The resolution of the final maps was estimated from the gold standard Fourier shell correlation (FSC = 0.143) criterion. Directional FSC plots were calculated on the 3DFSC server (54). The final cryo-EM map of the H3K56ac nucleosome was flipped along the z-axis using UCSF ChimeraX (55). Cryo-EM maps were low-pass filtered to 3.1 Å and normalized by root-mean-square scaling relative to the unmodified nucleosome map. All maps were displayed at the contour levels described in the figure legends.

### Model building and refinement

The initial atomic models were built based on the coordinates of the modified Widom 601 DNA sequence from the RNAPII-nucleosome complex (Protein Data Bank (PDB) ID: 6A5O (33)) and the coordinates of the histone octamer from the human nucleosome (PDB ID: 7VZ4 (56)). Acetyl-lysine residues were introduced using Coot (57). The atomic coordinates were fitted to the cryo-EM maps by rigid-body fitting using UCSF ChimeraX (55). The fitted coordinates were refined by phenix.real_space_refine (58), and manually adjusted by Coot (57) and ISOLDE (59). Refinement and validation statistics are summarized in Table 1. The distances between atoms were measured using USCF ChimeraX (55).

### Micrococcal nuclease assay

Nucleosomes (0.125 µM) were incubated in a reaction mixture, containing 10 mM HEPES-KOH (pH 7.5), 2.5% glycerol, 1.9 mM dithiothreitol, 40 mM Tris-HCl (pH 8.0), 25 mM sodium chloride, 2.5 mM calcium chloride, and 10 mU/µl micrococcal nuclease (Takara) at 37°C for 10 minutes. A 4 µl portion of the reaction mixture was mixed with 2 µl of deproteinization solution, containing 20 mM Tris-HCl (pH 8.0), 20 mM EDTA, 0.1% SDS, and 0.5 mg/ml proteinase K (Roche), and incubated at room temperature for 10 minutes. The samples were analyzed by 8% non-denaturing PAGE and stained with ethidium bromide. Gel images were acquired by an Amersham Imager 680 (Cytiva). The band intensities of the DNA were quantified using the ImageQuant™ TL software (GE Healthcare) and normalized with those of the undigested DNA. The assay was performed in technical triplicate, and the mean values are presented with standard deviations shown as error bars.

### Thermal stability assay

The thermal stability assay was conducted as reported previously, with slight modifications (60). The reaction buffer contained 0.125 µM nucleosome, 100 mM sodium chloride, 19 mM HEPES-KOH (pH 7.5), 0.95 mM DTT, 4.75% glycerol, and 1×protein thermal shift dye (Thermo Fisher Scientific). Samples were incubated in a QuantStudio 5 real-time PCR system (Thermo Fisher Scientific), and fluorescence was monitored during a temperature ramp from 25°C to 95°C. The temperature was increased in 1°C increments and maintained for 1 min at each step before fluorescence measurement. Fluorescence was detected using excitation filter 1 (470 nm) and emission filter 3 (587 nm). The fluorescence intensity at each temperature (F(T)) was normalized according to the following equation: (F(T) – F_25°C_) / (F_95°C_ – F_25°C_), where F_25°C_ and F_95°C_ represent the fluorescence intensities measured at 25°C and 95°C, respectively. The assay was performed in technical triplicate, and the mean values are presented with standard deviations (SD) shown as a shaded region.

### The use of Large Language Models

ChatGPT, a Large Language Model developed by OpenAI, was used in a strictly auxiliary role to support the writing process and graph generation. This included assistance with phrasing, grammar correction, and the generation of code examples in Python. The authors remain fully responsible for the content and conclusions of the manuscript.

## DATA AVAILABILITY

The cryo-EM maps of the unmodified, H3K56ac, H3K64ac, and H3K122ac nucleosomes were deposited to the Electron Microscopy Data Bank (EMDB) and the Protein Data Bank (PDB), respectively. The accession codes are as follows: unmodified nucleosome (EMD-66767 and PDB ID 9XDM), H3K56ac nucleosome (EMD-66768 and PDB ID 9XDN), H3K64ac nucleosome (EMD-66769 and PDB ID 9XDO), and H3K122ac nucleosome (EMD-66770 and PDB ID 9XDP). Half maps, primary reconstructed maps, atomic coordinates, FSC curve data, and validation reports are available through the corresponding EMDB and PDB entries. Uncropped gel images and quantification values are provided in Source Data 1-6.

## Acknowledgements

We thank Y. Takeda and M. Dacher (The University of Tokyo) for their assistance. We also thank M. Goto and M. Henmi (RIKEN) for their assistance with protein purification.

## Author contributions

Conceptualization: TO, TK, GH, YT, SSe, HK

Methodology: TO, SH, TK, SSu, KN, GH, SSe, YT, HK

Resources: TO, SSu, KN, GH, JK, YK, HE, HK

Investigation: TO, SH, SSu, KN, GH, YT, HK

Visualization: TO, SH, MO, YT, HK

Funding acquisition: TO, SH, TK, GH, SSe, YT, HK

Project administration: TK, GH, SSe, YT, HK

Supervision: TK, GH, SSe, YT, HK

Writing – original draft: TO, HK

Writing – review & editing: TO, SH, TK, GH, HE, SSe, YT, HK

## Funding and additional information

This work was supported in part by JSPS KAKENHI grant numbers JP25KJ1033 [to T.O.], JP25K18410 [to S.H.], JP23K17392 [to T.K.], JP24H00062 [to S.Se. and T.K.], JP26K01945 [to Y.T.], JP25K24594 [to H.K.], JP24H02319 [to H.K.], and JP24H02328 [to H.K.]; JST SPRING Grant Number JPMJSP2108 [to T.O.]; JST PRESTO Grant Number JPMJPR25N9 [to T.K.]; JST ERATO Grant Number JPMJER1901 [to H.K.]; JST CREST Grant Number JPMJCR24T3 [to H.K.]; KOSÉ Cosmetology Research Foundation [to Y.T.], Astellas Foundation for Research on Metabolic Disorders [to Y.T.], Nakatani Foundation [to H.K.]; and Research Support Project for Life Science and Drug Discovery (BINDS) grant numbers JP26ama121002 [to Y.T.] and JP26ama121009 [to H.K. and G.H.].

