## Supplementary Figure for "Structural basis for site-specific histone H3 acetylation-dependent regulation of RNAPII transcription through nucleosomes"

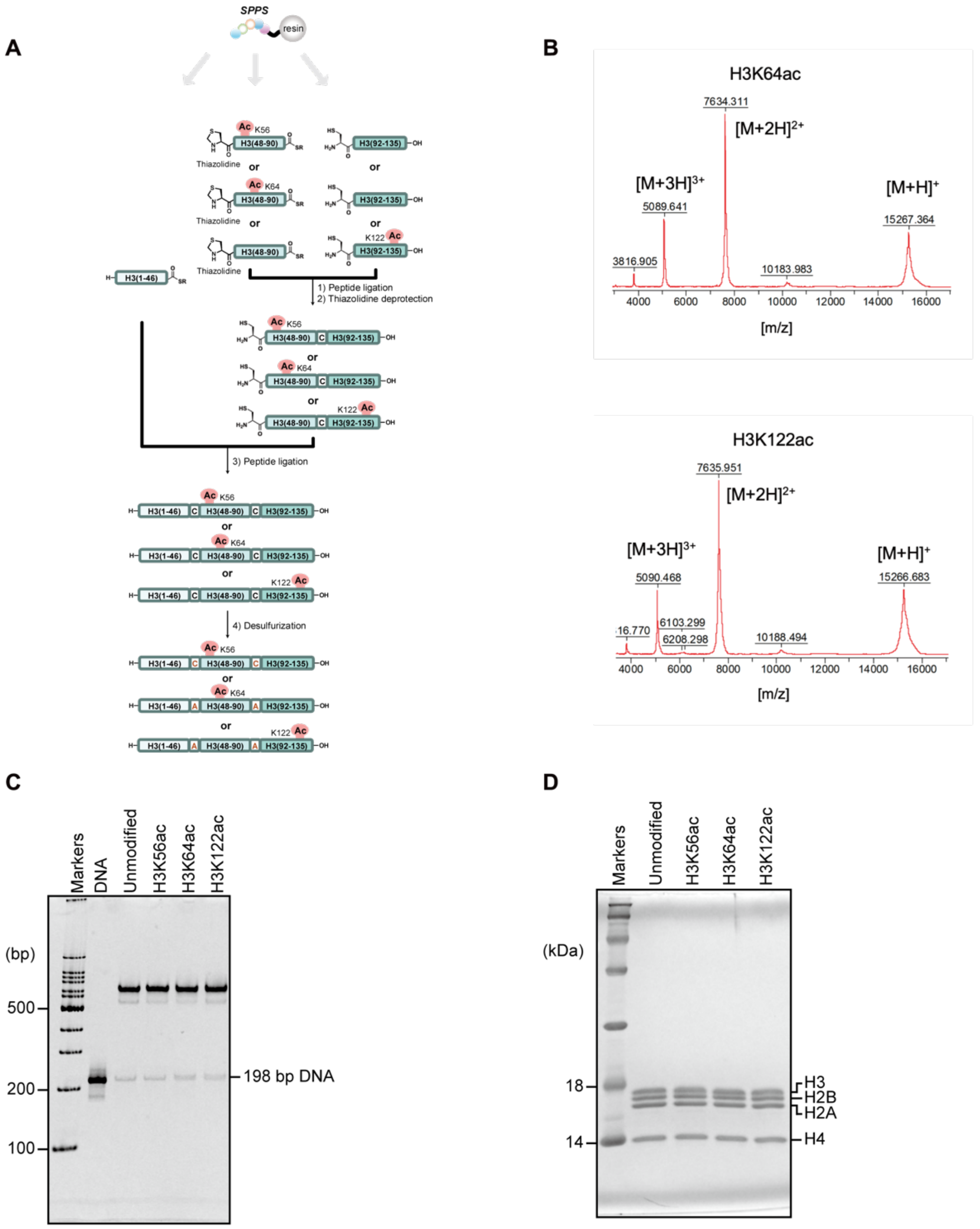


**Supplementary Figure 1: Preparation of acetylated nucleosomes.** (A) Schematic representation of chemical preparation of acetylated histones. (B) MALDI-TOF MS spectra of chemically synthesized H3K64ac and H3K122ac. m/z: [M+H]$^+$ Calcd. 15267.6; Found 15267.4 for H3K64ac and [M+H]$^+$ Calcd. 15267.6; Found 15266.7 for H3K122ac. (C) Free DNA and nucleosomes containing unmodified, H3K56ac, H3K64ac, and H3K122ac H3 peptides were analyzed by non-denaturing PAGE, with ethidium bromide staining. (D) Nucleosomes containing unmodified, H3K56ac, H3K64ac, and H3K122ac H3 peptides were analyzed by SDS-PAGE, with Coomassie Brilliant Blue staining.

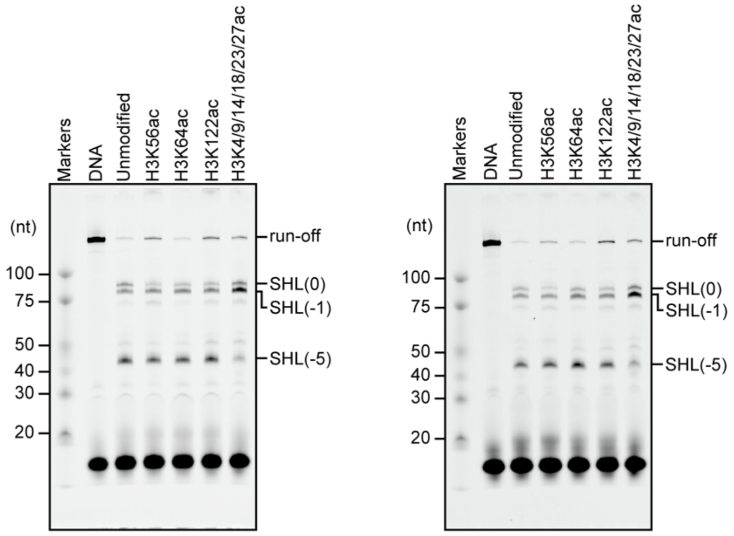


**Supplementary Figure 2: Repeated transcription assay of nucleosomes containing acetylated H3, as shown in Figure 1 (D).**

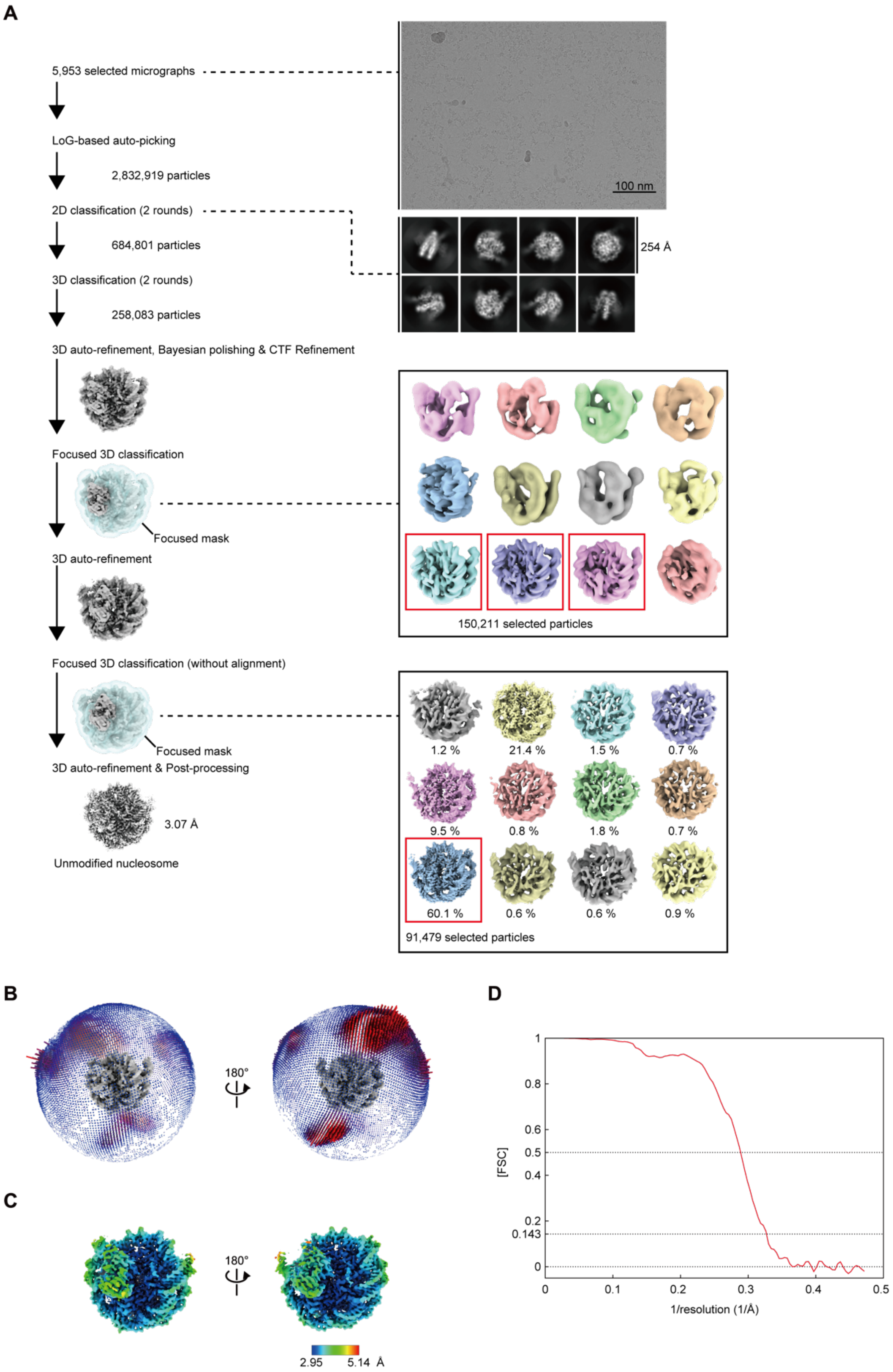


**Supplementary Figure 3: Cryo-EM analysis of the unmodified nucleosome.** (A) Image processing workflow of the unmodified nucleosome. (B) Euler angular distribution, (C) local resolution map, and (D) Fourier shell correlation (FSC) curve of the unmodified nucleosome.

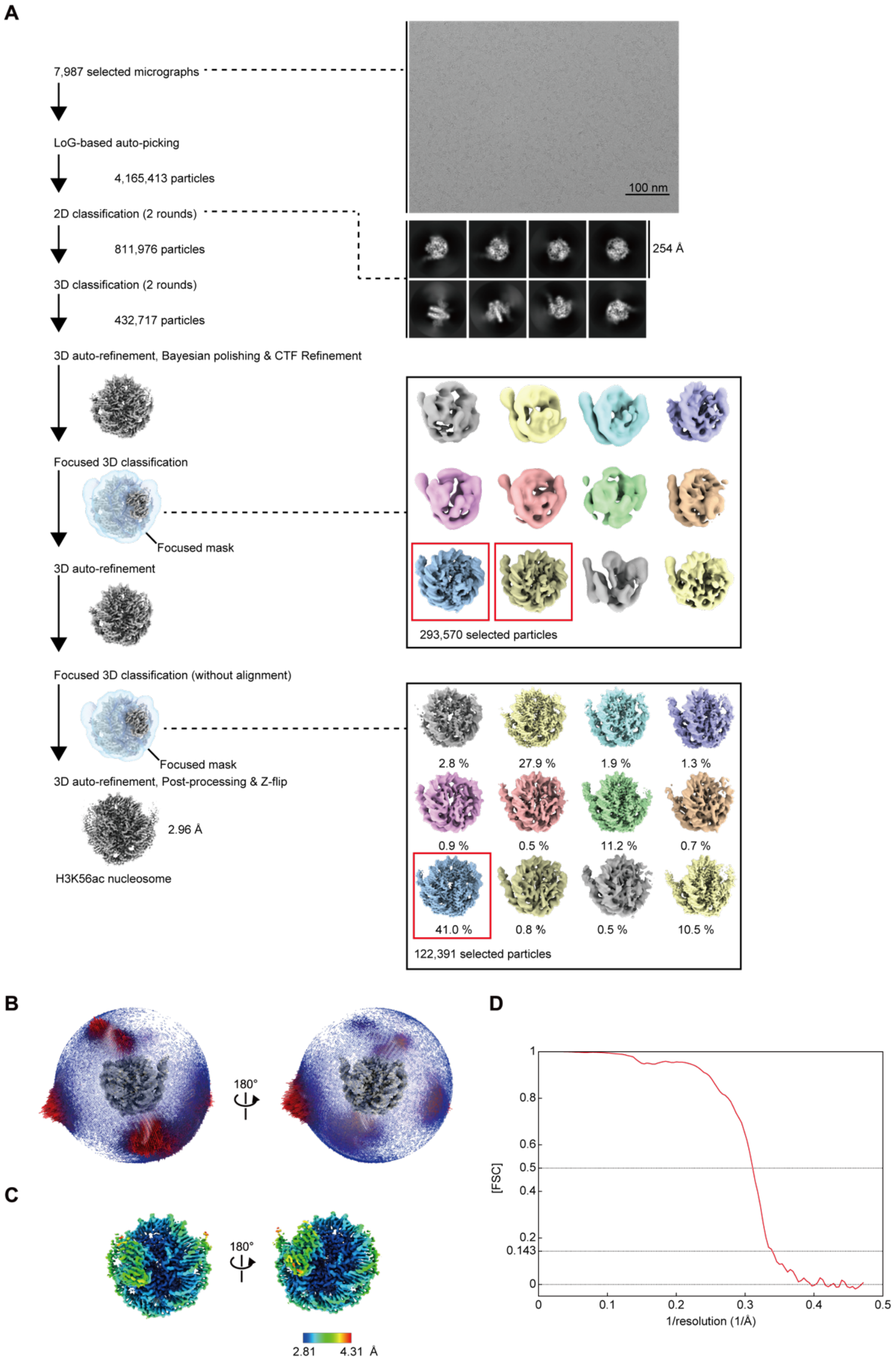


**Supplementary Figure 4: Cryo-EM analysis of the H3K56ac nucleosome.** (A) Image processing workflow of the H3K56ac nucleosome. (B) Euler angular distribution, (C) local resolution map, and (D) Fourier shell correlation (FSC) curve of the H3K56ac nucleosome.

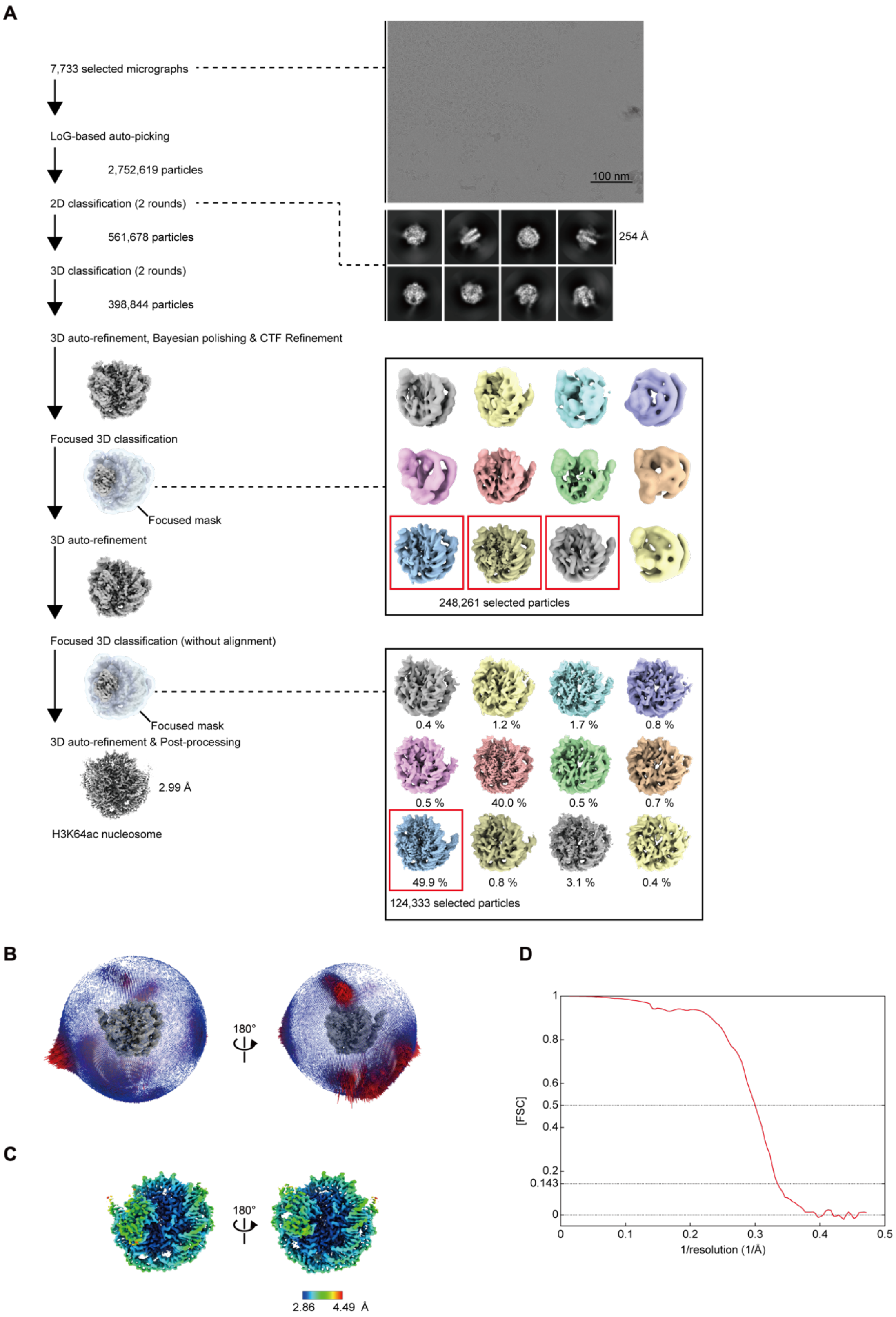


**Supplementary Figure 5: Cryo-EM analysis of the H3K64ac nucleosome.** (A) Image processing workflow of the H3K64ac nucleosome. (B) Euler angular distribution, (C) local resolution map, and (D) Fourier shell correlation (FSC) curve of the H3K64ac nucleosome.

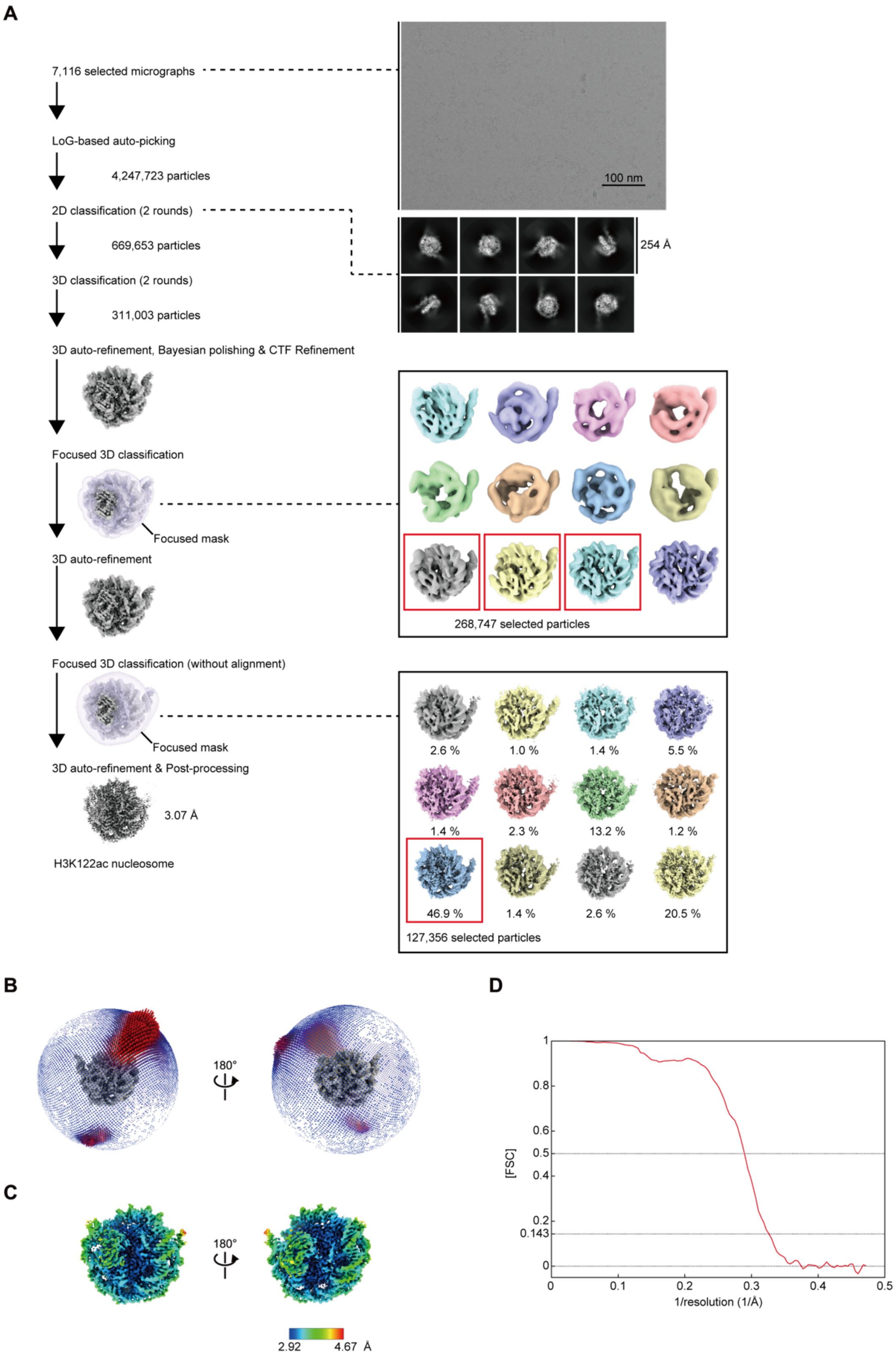


**Supplementary Figure 6: Cryo-EM analysis of the H3K122ac nucleosome.** (A) Image processing workflow of the H3K122ac nucleosome. (B) Euler angular distribution, (C) local resolution map, and (D) Fourier shell correlation (FSC) curve of the H3K122ac nucleosome.

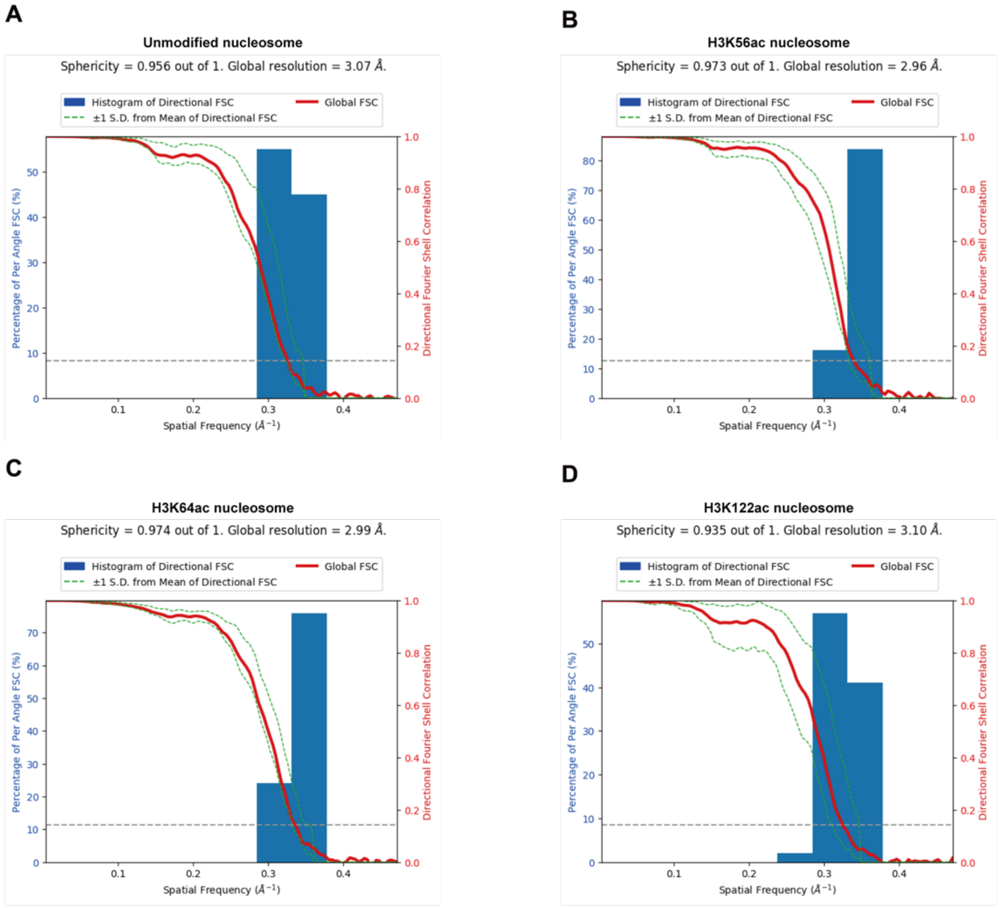


**Supplementary Figure 7. Directional FSC plots for the (A) unmodified, (B) H3K56ac, (C) H3K64ac, and (D) H3K122ac nucleosomes.**

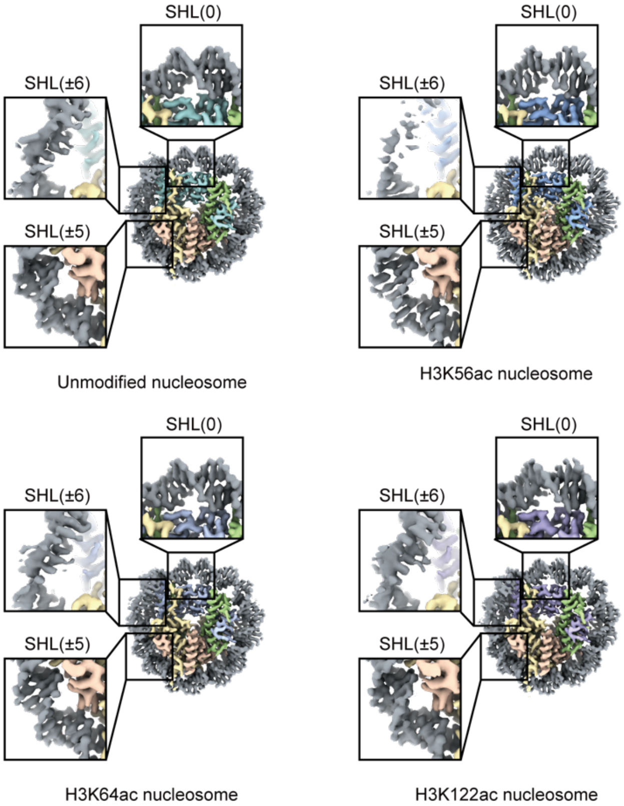


**Supplementary Figure 8. Close-up views of the DNA regions around SHL(±6), SHL(±5), and SHL(0).**

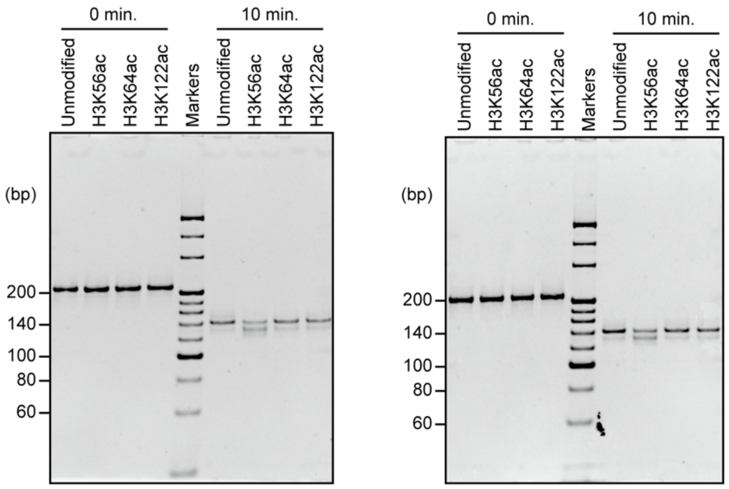


**Supplementary Figure 9: Repeated Micrococcal nuclease digestion assay of nucleosomes containing acetylated H3, as shown in Figure 3 (B).**

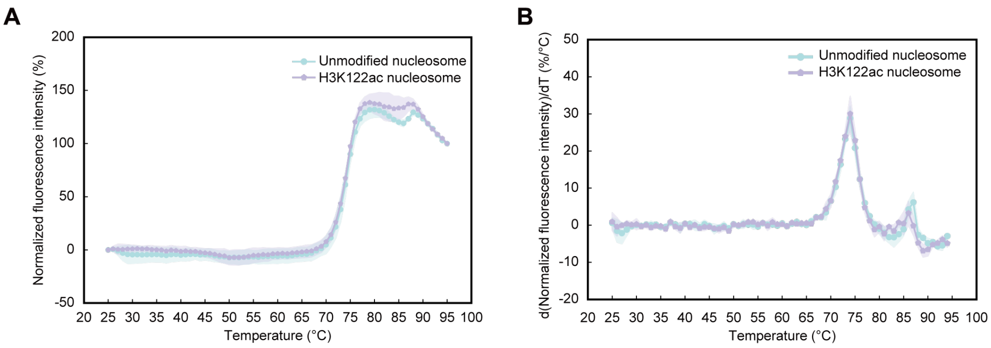


**Supplementary Figure 10. Thermal stability assay of the unmodified and H3K122ac nucleosomes.** (A) Normalized fluorescence intensity curves of the unmodified and H3K122ac nucleosomes. (B) Derivatives of the normalized fluorescence intensity curves shown in (A).
